# Differential cell fates of muscle stem cells are accompanied by symmetric segregation of canonical H3 histones *in vivo*

**DOI:** 10.1101/499913

**Authors:** Brendan Evano, Gilles Le Carrou, Geneviève Almouzni, Shahragim Tajbakhsh

## Abstract

Stem cells are maintained through symmetric or asymmetric cell divisions. While various mechanisms initiate asymmetric cell fates during mitosis, possible epigenetic control of this process has emerged recently. The asymmetrical distribution of a canonical histone H3 variant during mitosis in fly germline has suggested a role for partitioning old and new nucleosomes in asymmetric cell fates. Here, we provide resources for single cell assays and show the asymmetric segregation of transcription factors along with old and new DNA in mouse muscle stem cells *ex vivo* and *in vivo*. However, these differential fate outcomes contrast with a symmetric distribution of the canonical H3.1 vertebrate variant. These findings point to different evolutionary mechanisms operating in fly germline stem cells and vertebrate somatic stem cells to mitigate epigenetic regulation of asymmetric cell fates.

---

A long-standing question in multicellular organisms is how epigenetic information is transmitted or changed through cell division. The nucleosome is the basic unit of chromatin and notably contributes to gene expression regulation through its composition (histone variants and modifications). It is therefore particularly critical to determine how, and where in the genome, the choice of histone variants, their turnover and post-translational modifications contribute to the maintenance or switch of a particular cell fate.

Histone dynamics has been addressed with a variety of approaches, on bulk chromatin or at specific loci. Stable isotope labeling followed by mass spectrometry analysis notably revealed differences in turnover rates of the histone H3 variants H3.1 and H3.3^1^. Fluorescent tags were used to assess global incorporation or inheritance patterns of H3 variants in *D. melanogaster*^2,3^. Additionally, photo-bleaching, photo-activation and photo-conversion have been used to measure histone dynamics on short timescales^4^. Complementary approaches were developed to measure locus-specific histone dynamics, by sequential expression of tagged histones followed by chromatin immunoprecipitation and DNA sequencing^5,6^, which have poor temporal resolution, or by metabolic labeling of histone proteins, which has a better resolution but does not distinguish between different histone variants^7^. Although these methods provided significant insights into chromatin dynamics, they suffer from specific downsides, from lack of positional information to delays in expression of inducible reporters. In addition, current knowledge about histone dynamics arises predominantly from *in vitro* models where cell fate cannot be manipulated.

Histone H3 variants have a major role in epigenetic maintenance during development and disease^8^’^9^, but tools to measure histone dynamics *in vivo* either globally or at specific loci are lacking. Here, we sought to investigate histone H3 variants dynamics *in vivo* in mouse, using SNAP tags as reporters and skeletal muscle stem cells as a model of changes in cell fates. SNAP tags react specifically and irreversibly with benzylguanine derivatives^10^. A number of cell-permeable SNAP substrates are available (fluorescent, biotinylated or amenable to custom-probe synthesis through benzylguanine building blocks), with low toxicity and allowing fast and quantitative labeling^11^. SNAP tags were reported to work efficiently and specifically *in vivo* in mice^12^, and proved to be superior to classical fluorescent proteins in terms of spectral flexibility, signal-to-noise fluorescence and resistance to histological sample preparations^12,13^. Further, SNAP tags allow the development of functionalities *via* synthetic substrates that cannot be achieved with classical genetically encoded reporters^14^. Finally, SNAP tags have been used extensively to measure histone H3 variants dynamics on bulk chromatin^15–18^, and were adapted recently to measure specific histone variants turnover genome-wide^19^.

In most tissues, stem cells self-renew and give rise to daughter cells committed to differentiation, thereby ensuring both tissue homeostasis and maintenance of the stem cell pool^20^. This can be achieved through different modes of cell division, either symmetric (Symmetric Cell Division, SCD), self-renewing or differentiating, or asymmetric (Asymmetric Cell Division, ACD) divisions. Fine-tuning the balance between these different modes is critical for proper tissue development, homeostasis and regeneration, and to prevent tumorigenesis^20^.

A variety of subcellular constituents, including transcripts, organelles, centrosomes or sister chromatids can distribute asymmetrically between daughter cells during cell division^21^. A major challenge is to understand how their biased segregation dictates asymmetric cell fates^22^. The hypothesis of a role for replication-coupled nucleosome assembly in determining asymmetric cell fates in *C. elegans* nervous system has raised much interest^23^. Furthermore, recent studies reported non-equivalent inheritance of specific histone variants during ACD of *D. melanogaster* male germline stem cells (GSCs)^3,24^. These findings along with the importance of chromatin in cell plasticity^9^ raised the possibility of an epigenetic mechanism for maintaining the identity of self-renewed stem cells while resetting the identity of their differentiating sibling cells.

Here we sought to investigate whether the non-equivalent inheritance of specific histone variants is conserved in mammals, using mouse skeletal muscle stem cells as a model. Adult skeletal muscle satellite (stem) cells are mostly quiescent during homeostasis. Following muscle damage, they proliferate and differentiate into muscle fibres by cell-cell fusion^25,26^. A fraction of the population self-renews and reconstitutes a *de novo* satellite cell pool. This is accompanied by a temporal expression of cell fate markers, such as the transcription factors Pax7 (stem), Myod (commitment) and Myogenin (differentiation). Satellite cells can divide symmetrically and asymmetrically during muscle regeneration^27–29^, which is critical for proper muscle regeneration and maintenance of the satellite cell pool^27,30,31^.

## Results

To address whether the mechanism uncovered in Drosophila GSCs operates in a somatic mammalian stem cell, we developed a pulse-chase approach to follow histone variants using SNAP tags. SNAP-tagged histones have been used for addressing human H3.1 and H3.3 turnover during DNA replication and repair^16–18^. The asymmetric inheritance of H3, but not H3.3, reported in *D. melanogaster* GSCs^3^ prompted us to generate mouse reporter lines expressing SNAP-tagged H3.1 and H3.3 to follow their inheritance patterns during ACD of satellite cells.

*H3.1-SNAP* and *H3.3-SNAP* transgenes were expressed in several tissues (Supplementary Fig. 1a), at low levels (< 20%) compared to endogenous histone expression (Supplementary Fig. 1b). SNAP labeling of embryonic fibroblasts derived from reporter mice showed ubiquitous *H3.1-SNAP* expression, whereas *H3.3-SNAP* expression was lower and variegated (data not shown). Furthermore, as reported^16^ we find that newly deposited H3.1-SNAP histones overlapped with sites of DNA replication, while newly deposited H3.3-SNAP histones were uncoupled from DNA replication (Supplementary Fig. 1c). These results indicate that the *Tg:H3.1-SNAP* and *Tg:H3.3-SNAP* lines are faithful reporters of H3.1 and H3.3 turnover in mouse primary cells.

**Fig. 1.**
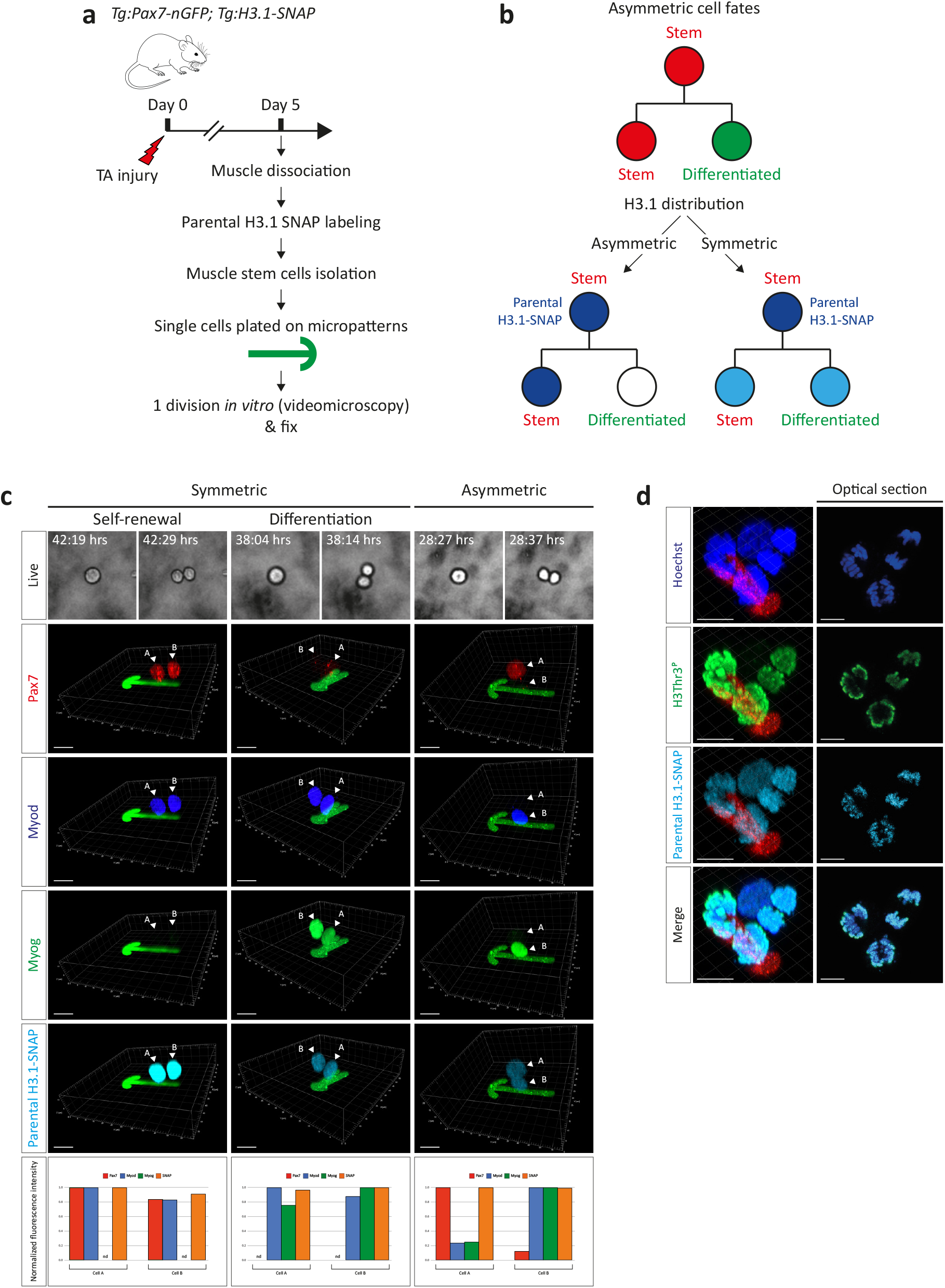
Parental H3.1 histones are symmetrically distributed during asymmetric cell fate decisions *ex vivo*. **a**, Experimental scheme. **b**, Potential parental histone distribution patterns. **s**, Representative examples of parental H3.1 inheritance pattern during SCD, towards selfrenewal (left) or differentiation (middle), or ACD (right). Arrowheads indicate nuclei of daughter cells. For each cell pair, the daughter cells were named ‘Cell A’ and ‘Cell B’. Fluorescence intensity was measured on segmented nuclei and expressed as a fraction of the maximum intensity between Cell A and Cell B. In case of absence of signal in both daughter cells (*eg*. symmetric absence of Pax7), the normalized fluorescence intensity was not determined (nd value). n=4 mice, n=100 cell divisions/mouse. **d**, H3T3^P^ signal colocalizes with H3.1-SNAP signal. n=3 mice. n=10 cell divisions/mouse. Scale bar 10 *μ*m.

To examine the distribution of parental H3.1 histones in satellite cells, adult *Tibialis anterior* (TA) muscles were injured to induce satellite cell activation and muscle regeneration. Parental H3.1 histones were SNAP-labeled prior to satellite cell isolation (labeling efficiency ≈ 80%, Fig. 1a and Supplementary Fig. 1d). Satellite cells were isolated from compound transgenic *Tg:Pax7-nGF*^32^ and *Tg:H3.1-SNAP* mice by fluorescence-activated cell sorting (FACS) where GFP intensity can be used to fractionate subpopulations with distinct properties, such as Pax7-nGFP^Hi^ cells previously reported to divide asymmetrically 5 days post-injury (dpi)^28^. *In vivo* activated/SNAP-labeled Pax7-nGFP^Hi^ cells were isolated by FACS and the ongoing cell division was monitored on micropatterns (Fig. 1a). Micropatterns provide a controlled microenvironment that allows confinement of single cells and their progeny^33^. We and others have shown that their geometry impacts on the frequency of ACD of stem cells^29,34^. If the parental histones are partitioned equally following mitosis, the fluorescent SNAP signal will be inherited equally between daughter cells. However, if the parental histones are inherited asymmetrically, the fluorescent SNAP signal will be detected only in a subset of daughter cells (Fig. 1b).

Modes of cell division were scored based on absolute presence or absence of cell fate markers. Their expression level and SNAP signal intensity were further measured quantitatively. We observed self-renewing and differentiating (Fig. 1c left and middle) SCDs, in addition to ACDs (Fig. 1c right), as reported^28,29^. Interestingly, we observed clones containing 4 cells with two self-renewed cells (Pax7^+^) and two differentiated cells (Myog^+^), indicating a switch from self-renewing SCD to ACD (Supplementary Fig. 1e). Among all observed clones, the SNAP signal intensity was equivalent between daughter cells, irrespective of their mode of cell division (Fig. 1c). The maximal SNAP intensity difference between daughter cells was less than 8% (Fig. 1c left), while transcription factors displayed differences in expression up to 80% between daughter cells in case of ACD (Fig. 1c right). These results indicate that parental H3.1 histones were essentially inherited symmetrically during both SCD and ACD of satellite cells.

H3-Thr3 phosphorylation (H3T3^P^) was reported to distinguish pre-existing and newly-deposited H3 in prophase, and to be required for asymmetric inheritance of old *vs*. new histones in *D. melanogaster* GSCs^24^. To further address the issue of histone inheritance, we investigated the distribution of H3T3^P^ in *in vivo* activated/SNAP-labeled Pax7-nGFP^Hi^ cells on micropatterns. We readily detected H3T3^P^ signals in mitotic cells, colocalizing with SNAP signals yet not restricted to a subset of chromatids in early mitosis (Fig. 1d). These observations indicate that H3T3^P^ did not point to a differential distribution of old *vs*. new histones in early mitosis of satellite cells.

Within adult muscles, satellite cells reside in a specific niche and a complex balance between extrinsic and intrinsic cues is required for proper satellite cell proliferation and cell fate determination^35^. As with other stem cell niches^36^, the composition of the satellite cell microenvironment is dynamic during muscle regeneration^37^, raising the possibility that some niche factors that cannot be modelled *ex vivo* might drive ACDs. Although ACDs have been reported extensively *ex vivo*, to the best of our knowledge, only one study has demonstrated asymmetry involving transcription factors in a vertebrate model system *in vivo*^38^.

Therefore, we established a clonal tracing strategy to examine two-cell clones in regenerating muscles *in vivo* (Fig. 2a and Fig. 2b), together with SNAP-labeling of parental H3.1 histones (*in vivo* labeling efficiency ≈ 60%, Supplementary Fig. 2b). As previously, modes of cell division were scored based on absolute presence or absence of cell fate markers. Their expression level and SNAP signal intensity were further measured quantitatively. We observed self-renewing (66.7%, n=98 of 147 cell pairs, Fig. 2c and Fig. 2d left) and differentiating (6.8%, n=10 of 147 cell pairs, Fig. 2c and Fig. 2d middle-left) SCDs, in addition to ACDs (23.1%, n=34 of 147 cell pairs, Fig. 2c and Fig. 2d middle-right and right). Notably, the relative frequencies of the different modes of cell division observed *in vivo* correspond to those on micropatterns *ex vivo*^29^. Among all clones analysed, we observed an equivalent SNAP signal between daughter cells (Fig. 2d). Interestingly, a difference in SNAP signal intensity (up to 40%) between daughter cells was noted for a limited number (≈ 5%) of divisions (Fig. 2d left and right, Fig. 2g right). However, the differences in SNAP intensity were lower than the observed differences in expression of cell fate markers, resulting from fluctuations in expression in case of SCD (Fig. 2d left) or asymmetric fate decisions (Fig. 2d right). Further, we did not detect examples showing exclusive distribution of parental histones to one daughter. These results point to a symmetric inheritance of parental H3.1 histones, irrespective of the mode of cell division *in vivo*.

**Fig. 2.**
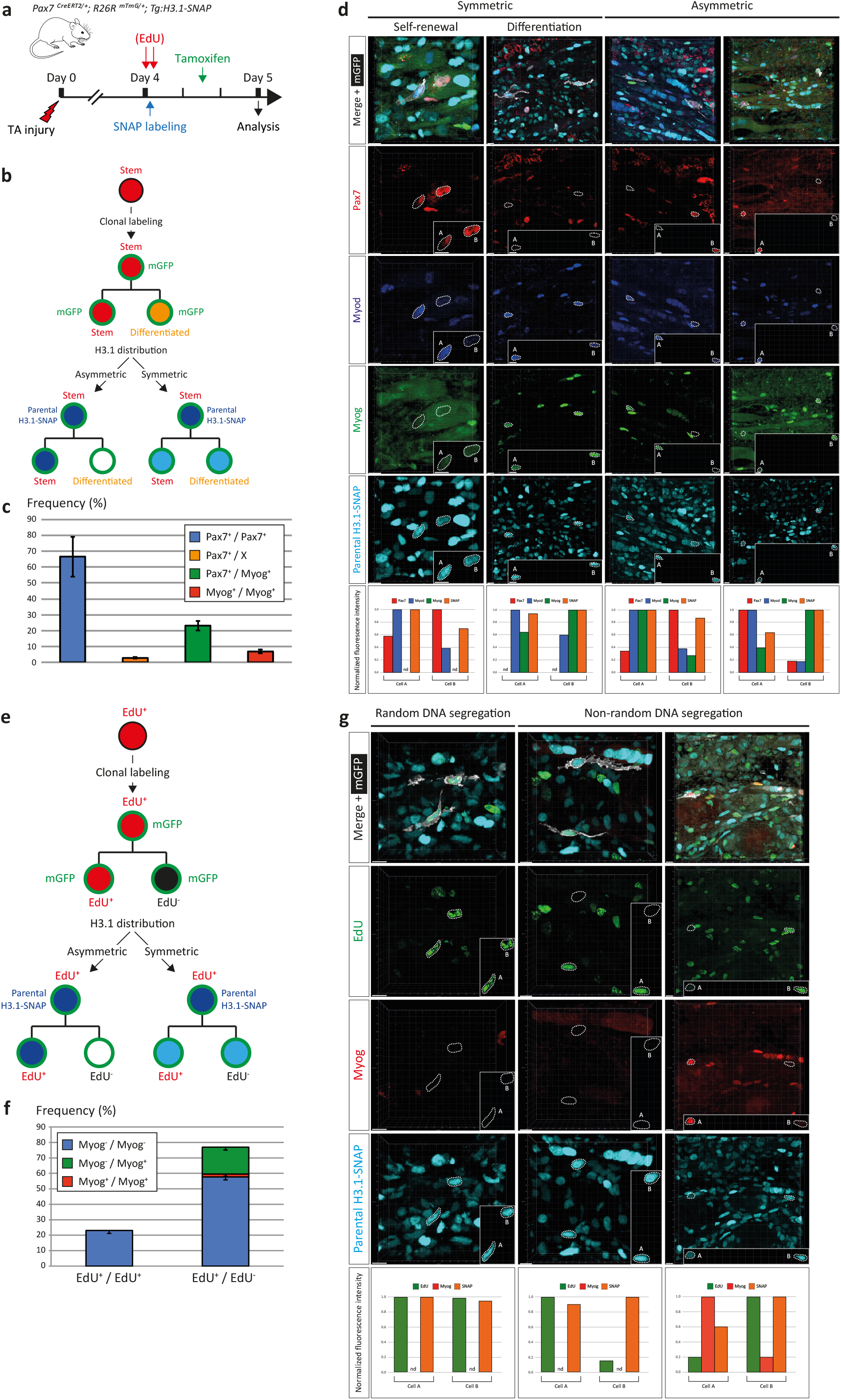
Parental H3.1 histones are symmetrically distributed during asymmetric cell fate decisions and non-random DNA segregation *in vivo*. **a**, Experimental scheme. Lineage-tracing was tamoxifen-induced 14 h before analysis; parental histones were labeled 8 h before tamoxifen administration; old DNA strands were labeled with EdU (Fig. 2e, 2f and 2g) 9 and 7 h before tamoxifen administration. **b**, Potential parental histone distribution patterns during asymmetric cell fate decisions. **c**, Frequencies of the different modes of cell division. Rare examples of Pax7/X and Myog/X daughter cells were observed, where one daughter cell expresses Pax7 or Myog and the other neither Pax7 nor Myog. n=3 mice, n=147 divisions. Data are presented as mean ± SEM. **d**, Representative examples of parental H3.1 inheritance pattern during SCD, towards self-renewal (left) or differentiation (middle-left), or ACD (middle-right and right). The mGFP signal from labeled daughter cells is shown only in the merged image for clarity; nuclei of daughter cells are outlined. Insets show nuclei of daughter cells isolated from other cells. For each cell pair, the daughter cells were named ‘Cell A’ and ‘Cell B’. Fluorescence intensity was measured on segmented nuclei and expressed as a fraction of the maximum intensity between Cell A and Cell B. In case of absence of signal in both daughter cells (*eg*. symmetric absence of Pax7), the normalized fluorescence intensity was not determined (nd value). See Supplementary Fig. 2a for corresponding low magnification images showing the clonality of the analysed events. Scale bar 10 *μ*m. **e**, Potential parental histone distribution patterns during non-random DNA segregation. **f**, Frequencies of the different modes of DNA segregation (random EdU^+^/EdU^+^ and non-random EdU^+^/EdU^-^) and fate decisions assessed by Myogenin expression. n=2 mice, n=52 divisions. Data are presented as mean ± SEM. **g**, Representative examples of parental H3.1 inheritance pattern during random DNA segregation (left) or non-random DNA segregation (middle and right), with symmetric absence of Myogenin expression (left and middle) or asymmetric Myogenin expression (right). The mGFP signal from labeled daughter cells is shown only in the merged image for clarity; nuclei of daughter cells are outlined. Insets show nuclei of daughter cells isolated from other cells. For each cell pair, the daughter cells were named ‘Cell A’ and ‘Cell B’. Fluorescence intensity was measured on segmented nuclei and expressed as a fraction of the maximum intensity between Cell A and Cell B. In case of absence of signal in both daughter cells (eg. symmetric absence of Myog), the normalized fluorescence intensity was not determined (nd value). See Supplementary Fig. 2c for corresponding low magnification images showing the clonality of the analysed events. Scale bar 10 *μ*m.

Notably, differences in SNAP signals between daughter cells were not observed *ex vivo* on micropatterns (Fig. 1c), suggesting variable H3.1 turnover rates between daughter cells *in vivo* and *ex vivo*. These observations highlight the importance of addressing epigenetic identity and plasticity *in vivo*. Finally, the SNAP signal was readily detected after histological preparation (strong fixation and heat-induced epitope retrieval), while the mGFP fluorescence was lost and required immunodetection, confirming the superiority of synthetic SNAP fluorophores over classical fluorescent proteins.

The asymmetric segregation of stem (Pax7) and differentiation (Myogenin) transcription factors was previously reported to correlate in part with non-random DNA segregation (NRDS) *ex vivo*, where old DNA strands are inherited by the stem cell and new DNA strands by the committed cell^28,29^. NRDS has been hypothesized to be a consequence of epigenetic mechanisms, possibly involving distribution of old and new histones or nucleosome pools^22^; however NRDS has not been shown robustly *in vivo* largely due to technical challenges. To examine parental H3.1 histone distribution and NRDS *in vivo*, we adopted the same strategy as above, together with a 5-ethynyl-2’-deoxyuridine (EdU) pulse-chase labeling (Fig. 2a and Fig. 2e) to identify both old DNA strands (EdU pulse) and new DNA strands (EdU chase).

We observed clonal examples of random DNA segregation (23.1%, n=12 of 52 cell pairs, Fig. 2f and Fig. 2g left) and non-random DNA segregation (76.9%, n=40 of 52 cell pairs, Fig. 2f and Fig. 2g middle and right). Interestingly, 22.5% of the cell pairs with NRDS showed asymmetric Myogenin expression (n=9 of 40 NRDS cell pairs, Fig. 2f and Fig. 2g right), with Myogenin being expressed in the EdU-negative daughter cell, as reported previously *ex vivo* where parental DNA strands are labeled and inherited by the Pax7-expressing cell^28,29^. Among all the clones analysed, the SNAP signal was equivalent between daughter cells (Fig. 2g), indicating a symmetric parental H3.1 inheritance, independent of the mode of DNA segregation.

## Discussion

Here we demonstrate that SNAP tags are powerful tools to investigate histone dynamics in mice *in vivo*. Our results indicate that parental H3.1 histones are equally inherited in the vast majority of dividing mouse satellite cells, *ex vivo* as *in vivo*, during asymmetric cell fate decisions with defined stem cell and differentiated cell transcription factors, as well as non-random segregation of DNA. Asymmetric histone H3 inheritance has uniquely been reported in *D. melanogaster* GSCs to date and was correlated with the execution of different cell fate programs following cell division^3,24^. A two-step mechanism was proposed where old and new histones are incorporated into different sister chromatids during S phase, followed by differential recognition and segregation of old and new sister chromatids by the mitotic apparatus^3,24^. This requires both an asymmetric distribution of histones at the replication fork, and strictly unidirectional DNA replication throughout the genome. Recently, two reports highlighted that while histone partitioning in the wake of the replication fork is inherently asymmetric between leading and lagging DNA strands, components of the replication machinery compensate to ensure an equal distribution of parental histones between sister chromatids^39,40^. Asymmetric histone distribution in Drosophila GSCs might then result from retention of parental histones on the leading strand and preferential unidirectional DNA replication^41^. While our results address global inheritance of parental histones and cannot exclude local limited asymmetry, notably at the replication fork, the discrepancy observed between *D. melanogaster* GSCs and our analysis of mouse satellite cells might indicate that the molecular mechanisms of asymmetric histone inheritance are not generally conserved through evolution, or alternatively between germline and somatic stem cells.

Our results show unambiguously for the first time NRDS *in vivo* and indicate that this process is not driven by a global asymmetric inheritance of old and new pools of H3.1. Nevertheless, according to the silent sister hypothesis^42^, epigenetic differences might exist between sister chromatids due to their relative difference in age. Asymmetric recognition of centromeres would then be the basis for NRDS, which would in turn lead to asymmetric gene expression and cell fate decisions. In this context, it is interesting to note that CENP-A, the centromeric H3 variant, was shown to be inherited asymmetrically during stem cell divisions in the fly midguts^43^, a possibility to explore in the future.

Finally, while our study investigated inheritance of bulk histones, the tools and methodologies reported here, combined with the recent advent of time-ChIP^19,44^, will allow genome-wide tracking of histone variant turnover rates. This can be done in a wide variety of mouse primary cell types in their *in vivo* environment thereby facilitating investigations of how epigenetic identity is maintained in stem cells, and remodeled in their differentiating progeny.

## Methods

### Mouse strains

*Tg:Pax7-nGFP* ^32^, *Pax7^CreERT2^* ^45^ and *R26^mTmG^* ^46^ were described previously. Breeding was performed on an F1:C57BL6/DBA2 background.6-8 weeks-old male littermates were used in this study, heterozygous for each allele used. Animals were handled according to national and European community guidelines, and protocols were approved by the ethics committee at Institut Pasteur.

### Cell lines and culture conditions

Primary fibroblasts were derived at E13.5 and cultured in DMEM GlutaMAX (ThermoFisher) 10% Fetal Bovine Serum (FBS) 1% Penicillin-Streptomycin (PS, ThermoFischer) at 37°C 5% CO_2_ 3% O_2_.

Satellite cells were plated on glass coverslips coated with Matrigel (Corning) for 30 min at 37°C or on micropatterns in satellite cell medium (40% DMEM 40% MCDB (Sigma) 20% FBS 2% Ultroser (Pall) 1% PS).

### Establishment of H3.1/H3.3-SNAP mouse strains

Mouse H3.1 (resp. H3.3) coding sequence was PCR amplified (Phusion Taq polymerase, New England Biolabs) from mouse primary fibroblasts cDNA (SuperScript III Reverse Transcriptase, ThermoFisher) with Notl_mH3.1_F (resp. Notl_mH3.3_F) and DPPVAT_mH3.1_R (resp. DPPVAT_mH3.3_R) primers, PCR-fused to a SNAP tag amplified from pSNAPm (New England Biolabs) with DPPVAT_SNAP_F and BamHI_SNAP_R primers, and cloned in pGEM-T easy (Promega) with T4 DNA ligase (New England Biolabs). A DPPVAT linker was included between histone coding sequences and the SNAP tag as in ^16,17^. H3.1-SNAP (resp. H3.3-SNAP) EcoRI/BamHI (New England Biolabs) fragment was subcloned in a pCAG SV40 Puro plasmid between the EcoRI and BamHI sites downstream the CAG promoter and upstream an SV40 early polyA termination sequence. CAGG H3.1/H3.3-SNAP polyA SpeI/KpnI (New England Biolabs) DNA fragments were microinjected in fertilized oocytes to derive transgenic lines.

Genotyping was performed by PCR from ear punch biopsies with CAG_For and DPPVAT_mH3.1_R or DPPVAT_mH3.R_R primers. *H3.1-SNAP* transgene insertion was mapped to chromosome 7 by inverse PCR (gDNA digestion with SacI (New England Biolabs), intramolecular ligation with T4 DNA ligase, PCR amplification with SV40 PolyA_F and SNAP qPCR_R primers). *H3.3-SNAP* transgene insertion was mapped to chromosome 10 by inverse PCR (gDNA digestion with BamHI, intramolecular ligation with T4 DNA ligase, PCR amplification with SNAP qPCR_F and CAG_R3 primers). Primers used are listed in Supplementary Table 1.

### RT-qPCR

Organs were homogenized in TRIzol (ThermoFisher) and RNA extracted following manufacturer’s instructions. gDNA was eliminated with Ambion DNAseI (ThermoFisher) and reverse transcription performed with SuperScript III Reverse Transcriptase (ThermoFisher). RNA was eliminated with RNAseH endonuclease (Roche) for 20 min at 37°C. mRNAs level was assessed with SYBR green master mix (Roche) and analysis was performed using the 2^-ΔCt^ method ^47^. RT-qPCR analyses have been normalized with *Tbp*. Specific forward and reverse primers used for RT-qPCR are listed in Supplementary Table 1.

### Western blot

Histones were extracted following ^48^ and protein concentration measured with the Pierce BCA Protein Assay Kit (ThermoFisher). 1 *μ*g of histones was run on 4-12% Bis-Tris Gel NuPAGE (ThermoFisher) in NuPAGE MES SDS Running Buffer (ThermoFisher) and transferred on a nitrocellulose membrane (Hybond ECL, Sigma) in 1X Tris-Glycine-SDS buffer (Euromedex) with 20% ethanol. Equivalent loading was evaluated from Ponceau staining of the membrane. The membrane was then blocked with 5% non-fat milk in Tris Buffer Saline (TBS) Tween 0.05% (TBS-T) for 20 min at RT and probed with specific primary antibodies overnight at 4°C in TBS-T 5% milk. After three washes in TBS-T, the membrane was incubated with HRP-conjugated secondary antibodies in TBS-T for 30 min at RT, washed three times in TBS-T, and revealed by chemiluminescence (Pierce ECL Plus Western Blotting Substrate, ThermoFisher) with autoradiography films (Hyperfilm ECL, GE Healthcare).

### SNAP labeling

*In vitro* SNAP quenching (Supplementary Fig. 1c) was performed with 10 μM SNAP Cell Block (New England Biolabs) for 30 min at 37°C, followed by three washes with culture medium. *In vitro* SNAP labeling was performed with 3 *μ*M SNAP Cell TMR Star (New England Biolabs) in culture medium for 20 min at 37°C, followed by three washes with culture medium, and washed in culture medium for 30 min at 37°C. Sites of DNA replication were labeled with 1 *μ*M EdU (ThermoFisher) for 10 min at 37°C. Soluble histones were extracted with CSK buffer prior to fixation as in ^16^. EdU incorporation was detected with the Click-iT Plus Alexa Fluor 488 Imaging Kit (ThermoFisher) as recommended.

*Ex vivo* SNAP labeling (Fig. 1a) was performed by incubating dissected and minced TA muscles in 1 ml satellite cell medium containing 3 *μ*M SNAP Cell SiR 647 (New England Biolabs, S9102S) for 30 min at 37°C with gentle agitation. Muscles were centrifuged at 500g for 5 min, supernatant was discarded and muscles resuspended in 1 ml DMEM. Muscles were centrifuged at 500g for 5 min, supernatant was discarded and muscles resuspended in collagenase D/tryspin mix for satellite cell isolation (see below).

*In vivo* SNAP labeling was performed by injecting 10 *μ*l of 30 *μ*M SNAP Cell SiR 647 (New England Biolabs) in NaCl 0.9% per TA at 4 dpi under isofluorane anesthesia, 8 h before tamoxifen administration (Fig. 2a). *In vivo* H3.1-SNAP labeling at 4 dpi enables specific labeling of ≈ 60% of satellite cells as early as 2 h post SNAP substrate injection (Supplementary Fig. 2b).

### Muscle injury, tamoxifen and EdU delivery

Muscle injury was done as described previously ^49^. Briefly, mice were anesthetized with 0.5% Imalgene/2% Rompun. The *Tibialis anterior* (TA) muscle was injected with 15 *μ*l of 10 *μ*M notexin (Latoxan) in NaCl 0.9%.

Tamoxifen (Sigma) was reconstituted at 25 mg/ml in corn oil/ethanol (5%) and stored at -20°C. Before use, tamoxifen was diluted at appropriate concentration (1 mg/ml or 0.25 mg/ml) with corn oil, and administered (8 *μ*l per g of mouse) by intragastric injection at 4 days post-injury and 14 h prior to sacrifice to allow for a maximum of two consecutive cell divisions *in vivo* (*in vivo* satellite cells doubling time about 8 h ^28^). We optimized conditions for clonal tracing leading to two-cell clones shortly after (14 h) tamoxifen administration. Clones were defined when mGFP-labeled cells were in close proximity (< 150 *μ*m apart, Supplementary Fig. 2a and Supplementary Fig. 2c) and isolated from any other mGFP-labeled cell. In addition, the definition of criteria for clonality was based on an extremely low dose of tamoxifen (2 *μ*g/g of mouse, recombination frequency < 0.5%). We then used a tamoxifen dose (8 *μ*g/g of mouse) inducing a recombination frequency about 10% to increase the representation of the myogenic population. For each mouse, one TA muscle was used for immunohistochemistry analysis, and the contralateral TA was used for determining the recombination efficiency following dissociation and immunostaining.

EdU (ThermoFisher) was dissolved in NaCl 0.9% and injected intraperitoneally (0.3 *μ*g/g of mouse) 9 and 7 h before tamoxifen administration.

### Satellite cells isolation and live-imaging

Dissections were done essentially as described previously ^49^. Injured TA muscles were removed from the bone in cold DMEM, minced with scissors, and then digested with a mixture of 0.08% Collagenase D (Sigma), 0.1% Trypsin (ThermoFisher), 10 *μ*g/ml DNase I (Sigma) in DMEM for five consecutive cycles of 30 min at 37°C. For each round, the supernatant was filtered through a 70 *μ*m cell strainer (Miltenyi) and collagenase and trypsin were blocked with 8% FBS on ice. Pooled supernatants from each digestion cycle were centrifuged at 500g for 15 min at 4°C. The pellet was resuspended in 2 ml cold PBS (ThermoFisher) and placed on top of cold Percoll layers (Sigma)(5 ml of 20% Percoll and 3 ml of 60% Percoll). After centrifugation at 500g for 15 min at 4°C, cells were collected from the 20%-60% interphase, while dense debris were concentrated below the 60% layer. The collected fraction was diluted in 40 ml DMEM 8% FBS and centrifuged at 500g for 15 min at 4°C. The pellet was resuspended in 500 *μ*l satellite cell medium and filtered through a 40 *μ*m cell strainer (Falcon). Cells were plated on glass coverslips coated with Matrigel or sorted (Supplementary Fig. 3) using a FACS Aria III (BD Biosciences) and collected in cold 250 μl of satellite cell medium. Sorted cells were plated on micropatterns (see below) and incubated at 37°C, 5% CO_2_, and 3% O_2_ in a Zeiss Observer.Z1 microscope connected to an LCI PlnN 10x/0.8 W DICII objective and Hamamatsu Orca Flash 4 camera piloted with Zen (Zeiss). Cells were filmed and images were taken every 9 min with bright-field filter to ensure that single cells were plated on the micropatterns and that their progeny remained confined. The raw data were analysed with TrackMate ^50^, transformed and presented as videos.

### Micropatterns

Micropatterns were manufactured essentially as described ^29,51^. Briefly, glass coverslips (24 × 60 mm #1, Thermo Scientific Menzel) were cleaned serially in H_2_O/acetone/isopropanol, activated with an oxygen plasma treatment (Harrick Plasma) for 5 min at 30W and incubated with poly-L-lysine polyethylene glycol (PLL(20)-g[3.5]-PEG(2), SuSoS) at 100 *μ*g/ml in 10 mM HEPES pH 7.4 at room temperature (RT) for 30 min, washed twice with H_2_0 and stored dried at 4°C. PLL-PEG-coated slides were placed in contact with an optical mask containing transparent micropatterns (Toppan Photomask) using H_2_O (0.6 *μ*l/cm^2^) to ensure tight contact between the mask and the coverslip and then exposed to deep UV light (Jelight). Micropatterned slides were subsequently incubated with 40 μg/ml fibronectin (Sigma) and 5 *μ*g/ml fibrinogen Alexa Fluor 488 (ThermoFisher) in 100 mM NaHCO_3_pH 8.3 for 25 min at RT and rinsed three times in NaHCO_3_, three times in H_2_0 and dried and transferred immediately on a 12 kPa polyacrylamide (PAA) gel on silanized glass coverslips, to match the substrate rigidity of skeletal muscles ^52^. 81 mm^2^-square glass coverslips were cut out larger coverslips (22 × 22 mm #1, Thermo Scientific Menzel), cleaned serially with H_2_O/EtOH 70%/Et0H 100%, deep-UV-activated, silanized with 7.1% (vol/vol) bind-silane (Sigma) and 7.1% (vol/vol) acetic acid in EtOH 99% for 10 min at RT, washed twice with EtOH 99% and dried. Transfer of micropatterns on PAA gel was performed with 7 *μ*l/cm^2^of a solution containing 7.5% acrylamide (Sigma), 0.16% bis-acrylamide (Sigma), 0.05% TEMED (Sigma), 0.05% APS (Sigma, A3678) in HEPES 10 mM pH 7.4 for 45 min at RT. Gels were rehydrated in NaHCO_3_ 100 mM pH 8.3 for 15 min at RT, detached from patterned coverslips and washed with PBS three times before cell seeding.

Micropatterned PAA gels were transferred in an in-house-made glass-bottom 6 well Petri dish to allow live-imaging and PAA embedding before immunostaining (see below). Briefly, 32 mm diameter glass coverslips were cleaned and silanized, a 14 × 14 × 0.45 mm piece of silicone (Smooth-on) was put at the center, and coverslips were layered with a 12 kPa PAA gel (‘surrounding’ PAA). After PAA polymerization, the coverslips were sealed (silicone) to the bottom of a 6 well plate (TPP) drilled out to 30 mm diameter. The silicone at the center of the PAA gel was removed and replaced with a 16 × 16 × 3 mm silicone isolator with a 10 × 10 × 3 mm window at the center to fit micropatterned PAA gels (9 × 9 mm) after filling with PBS. The apparatus was washed several times and equilibrated with satellite cell medium. Medium was removed completely and cells (250 *μ*l) seeded within the silicone isolator on micropatterns and, 90 min after plating, nonattached cells were washed with medium and medium volume was adjusted to 5 ml.

### Immunocytochemistry and Immunohistochemistry

Cells on micropatterns were fixed in 4% paraformaldehyde (PFA, Electron Microscopy Sciences) in PBS containing 0.9 mM CaCl_2_ and 0.5 mM MgCl_2_(PBS Ca^2+^ Mg^2+^) for 5 min at RT and washed in PBS Ca^2+^ Mg^2+^ for 5 min at RT. To prevent cell loss during subsequent immunostaining steps, cells were embedded in PAA. Cells were equilibrated in a solution containing 3.82% acrylamide, 0.13% bis-acrylamide and 0.1% TEMED in PBS for 30 min at RT, and the solution was replaced with 3.82% acrylamide, 0.13% bis-acrylamide, 0.1% TEMED and 0.05% APS in PBS. The silicone isolator was carefully detached and a parafilm-covered 20 × 20 mm glass coverslip (20 × 20 mm #1, Thermo Scientific Menzel) was adjusted on top of the micropatterns, sitting on the ‘surrounding’ PAA. Excess polymerization solution was removed and PAA allowed to polymerize for 1 h at RT. After rehydration in PBS for 15 min, the parafilm-covered coverslip was detached, PAA-embedded micropatterns were recovered and washed several times in PBS. Cells were permeabilized in 0.5% Triton X-100 (Sigma) for 20 min at RT, washed three times 10 min in PBS, blocked in 10% goat serum (GS, Gibco) for 4 h at RT, incubated with primary antibodies overnight at 4°C in 2% GS, washed four times 1 h in PBS at 4°C, incubated with Alexa Fluor-conjugated secondary antibodies and Hoechst when required overnight at 4°C in 2% GS and washed four times 1 h in PBS at 4°C. For Pax7/Myod/Myog triple staining, cells were further blocked with Pax7 primary antibody overnight at 4°C in 2% GS, washed four times 1 h in PBS at 4°C, incubated with Alexa Fluor 488-conjugated anti-Myogenin and DyLight 405-conjugated anti-Myod antibodies (see below) in 2% GS overnight at 4°C and washed four times 1 h in PBS at 4°C.

Cells on glass coverslips were fixed in 4% PFA for 5 min at RT, washed twice in PBS, permeabilized in 0.5% Triton X-100 for 5 min, blocked in 10% GS for 30 min at RT, incubated with primary antibodies in 2% GS for 2 h at RT, washed in PBS three times 5 min, incubated with Alexa Fluor-conjugated secondary antibodies and Hoechst for 45 min at RT, washed in PBS three times 5 min and mounted in PBS Glycerol 75%.

For immunohistochemistry, TA muscles were fixed for 24 h in 4% PFA at 4°C, washed in cold PBS four times 1 h at 4°C, equilibrated in 20 % sucrose (Sigma) overnight at 4°C, embedded in Tissue Freezing Medium (Leica) and snap-frozen on liquid nitrogen. 12 μm cryosections were allowed to dry at RT for 30 min and rehydrated in PBS for 15 min. Antigen retrieval was performed in Tris 10 mM EDTA 1 mM pH 9 for 10 min at 95°C, followed by three short PBS washes and an additional wash in PBS for 30 min at RT. Sections were permeabilized in 0.5% Triton X-100 for 20 min at RT, washed three times in PBS, blocked in M.O.M. Mouse Ig Blocking Reagent (Vector Laboratories) for 1 h at RT, washed three times in PBS, incubated in M.O.M. Diluent (Vector Laboratories) for 5 min at RT, incubated with primary antibodies (Pax7 (Fig. 2d) or 5FD (Fig. 2g), GFP, in M.O.M. Diluent) for 1 h at RT, washed three times 5 min in PBS, incubated with M.O.M. Biotinylated Anti-Mouse IgG Reagent (Vector Laboratories) for 10 min at RT, washed three times 5 min in PBS, incubated with secondary antibodies (Streptavidin-Alexa Fluor 405, Donkey anti-Chick Alexa Fluor 594, in M.O.M. Diluent) for 30 min at RT, washed three times 5 min in PBS. For Pax7/Myod/Myog triple staining, sections were further blocked with anti-Pax7 antibody (in M.O.M. Diluent) for 1 h at RT, washed three times 5 min in PBS, incubated with Alexa Fluor 488-conjugated anti-Myogenin and DyLight 550-conjugated anti-Myod antibodies (see below, in M.O.M. Diluent) overnight at 4°C, washed in PBS three times 5 min and mounted in PBS Glycerol 75%. EdU incorporation was detected with the Click-iT Plus Alexa Fluor 488 Imaging Kit (ThermoFisher) as recommended.

Immunostainings were analysed with a Zeiss Observer.Z1 and a Zeiss LSM800 confocal microscopes. Images were processed with Imaris 7.2.1 software. For each cell pair, the daughter cells were named ‘Cell A’ and ‘Cell B’. Fluorescence intensity was measured on segmented nuclei and expressed as a fraction of the maximum intensity between Cell A and Cell B. In case of absence of signal in both daughter cells (*eg*. symmetric absence of Pax7), the normalized fluorescence intensity was not determined (nd value).

### Antibody fluorescent labeling

Anti-myogenin antibody (5FD, DSHB) was purified with protein G sepharose (Sigma) as recommended. Briefly, hybridoma supernatant was incubated with protein G sepharose overnight at 4°C and packed into polypropylene columns (Qiagen), washed five times with PBS and antibody was eluted twice with 1.5 ml glycine 100 mM pH 2.7, neutralized with 150 *μ*l Tris 1M pH 9. Elution fractions were pooled and dialyzed (ThermoFisher) against PBS overnight at 4°C. Antibody was concentrated with Amicon Ultra centrifugal filters (Merck) to 1 mg/ml. 100 *μ*l of purified F5D antibody at 1 mg/ml (resp. anti-Myod at 0.75 mg/ml (BD Bioscience)) were dialysed (Thermofisher) against NaHCO_3_100 mM pH 8.3 (resp. Na_2_B_4_O_7_ 50 mM pH 8.5) overnight at 4°C, incubated with 120 *μ*M Alexa Fluor 488 NHS ester (ThermoFisher)(resp. 60 μM DyLight 405 NHS ester (ThermoFisher) or 70 *μ*M DyLight 550 NHS ester (ThermoFisher)) at 23°C for 1 h. NaCl concentration was adjusted to 150 mM and excess fluorescent dye was removed with Pierce Dye Removal Columns (ThermoFisher) following manufacturer’s recommendations. Desalted labeled antibodies were further dialysed (ThermoFisher, Cat#69570) against PBS overnight at 4°C and stored at -20°C with 1 mg/ml bovine serum albumin (Sigma) and 50% glycerol.

## Supporting information

## Data availability

All data supporting the findings of this study are available from the corresponding author on reasonable request.

## Acknowlegments

This work was supported by Institut Pasteur and grants from Agence Nationale de la Recherche (Laboratoire d’Excellence Revive, Investissement d’Avenir; ANR-10-LABX-73, ANR-16-CE12-0024) and Association Française contre les Myopathies, CNRS, and the European Research Council (Advanced Research Grant 332893). We would like to thank the Mouse Genetics Engineering Platform (Institut Pasteur), the Flow Cytometry and Photonic Bioimaging Platforms of the Center for Technological Resources and Research (Institut Pasteur), Johan van der Vlag (Radboud University, Netherlands) for providing the H3.1 antibody, Glenda Comai (Institut Pasteur) for initial assistance with confocal imaging, Dan Filipescu for his advice to establish the mouse H3-SNAP variants, and Tom Cheung (Hong Kong University of Science & Technology) for advice on antibody labeling. G.A. received support from ANR.

## Author contributions

B. E. and S.T. proposed the concept, designed the experiments, and wrote the paper. B.E. performed and analysed the experiments. G.L.C. provided technical support. G.A. contributed to application for the project, advised on experiments and edited the paper. All authors read and agreed on the manuscript.

## Competing interests

The authors declare no competing interests.

## Materials and correspondence

Correspondence and material requests should be addressed to

## Statistical information

Bar charts represent the mean ± standard error of the mean (SEM) or standard deviation of the mean (SD) as specified.

## Supplementary information

### Supplementary figures

**Supplementary Fig. 1.**
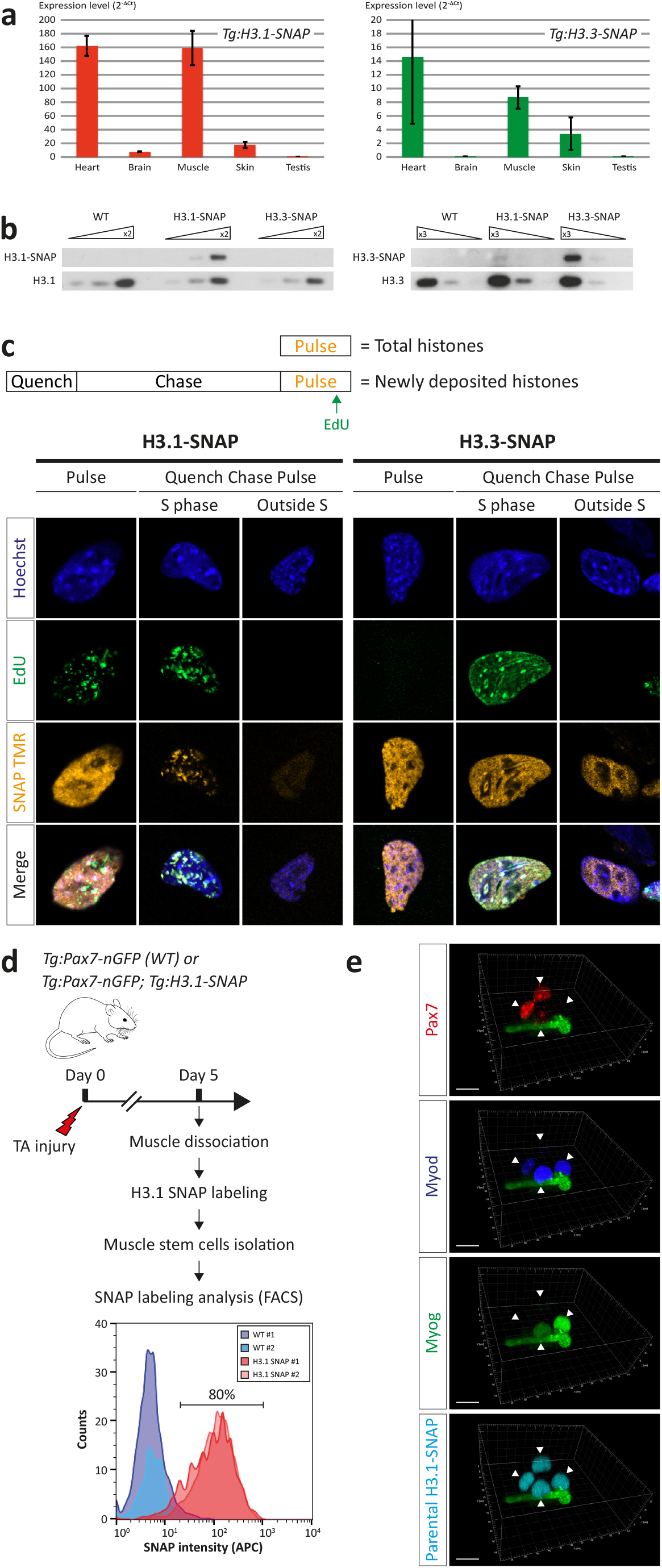
*Tg:H3.1-SNAP* and *Tg:H3.3-SNAP* reporter lines validation and *ex vivo* SNAP labeling of satellite cells. **a**, Transgene expression level (RT-qPCR) from heart, brain, skeletal muscle, skin and testis; n=2 mice per genotype. Data are presented as mean ± SD. **b**, Transgene expression level (Western blot) from primary embryonic fibroblasts derived from WT, *Tg:H3.1-SNAP* and *Tg:H3.3-SNAP mice*, detected with anti-H3.1 (left) or anti-H3.3 (right) antibodies. Each panel shows the endogenous (bottom) and SNAP-tagged (top) histone variant. Triangles indicate different amounts of material loaded, centered on 1 μg. **c**, SNAP TMR labeling of total (Pulse) or newly deposited (Quench Chase Pulse) SNAP-tagged histone variants and DNA replication sites (EdU) in *Tg:H3.1-SNAP* and *Tg:H3.3-SNAP* primary fibroblasts as in^16^. *d*, Efficiency of *ex vivo* SNAP labeling of satellite cells. 80% of *Tg:H3.1-SNAP* satellite cells were labeled compared to WT control cells; n=2 mice per genotype. **e**, Example of a four-cell clone with two Pax7+ cells and two Myog+ cells on a micropattern. Myog+ differentiated cells have left the cell cycle, therefore the 4-cell clone can result from an ACD where the Pax7+ daughter undergoes a SCD, then ACD; another possibility is an initial SCD, then each daughter undergoes an ACD. No cell death was observed. Scale bar 10 *μ*m.

**Supplementary Fig. 2.**
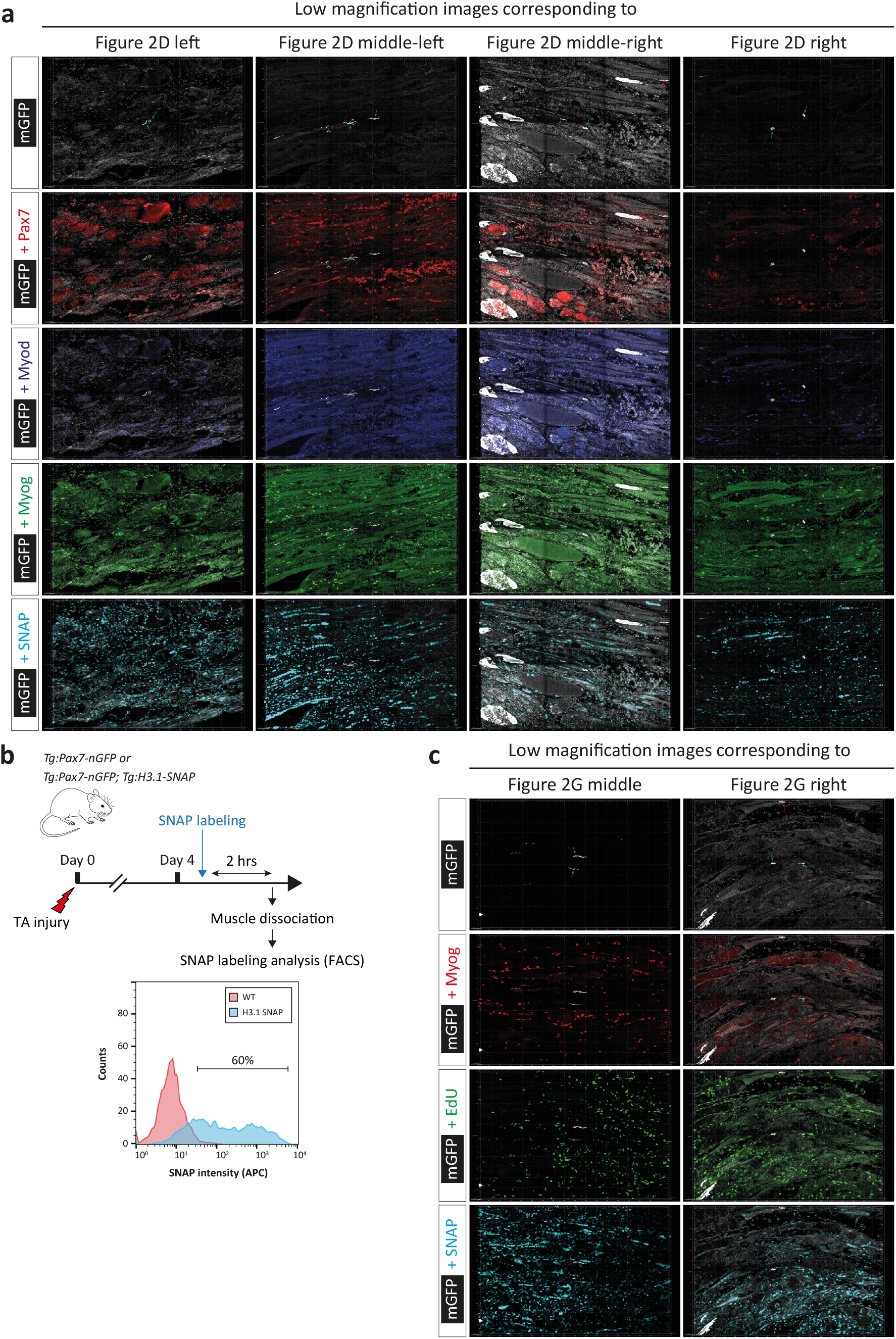
*In vivo* SNAP labeling of satellite cells and clonality of the analysed *in vivo* cell divisions. **a**, Low magnification images (600 *μ*m × 900 *μ*m) corresponding to high magnification images of Fig. 2d. Green arrows indicate mGFP-labeled daughter cells; red arrows indicate presumably unrelated mGFP-labeled cells. Scale bar 50 *μ*m. **b**, Efficiency of *in vivo* SNAP labeling of satellite cells at 4 dpi. 60% of *Tg:H3.1-SNAP* satellite cells were labeled compared to WT control cells; n=2 mice per genotype. c, Low magnification images (600 *μ*m × 900 *μ*m) corresponding to high magnification images of Fig. 2g. Green arrows indicate mGFP-labeled daughter cells; red arrows indicate presumably unrelated mGFP-labeled cells. Scale bar 50 *μ*m.

**Supplementary Fig. 3.**
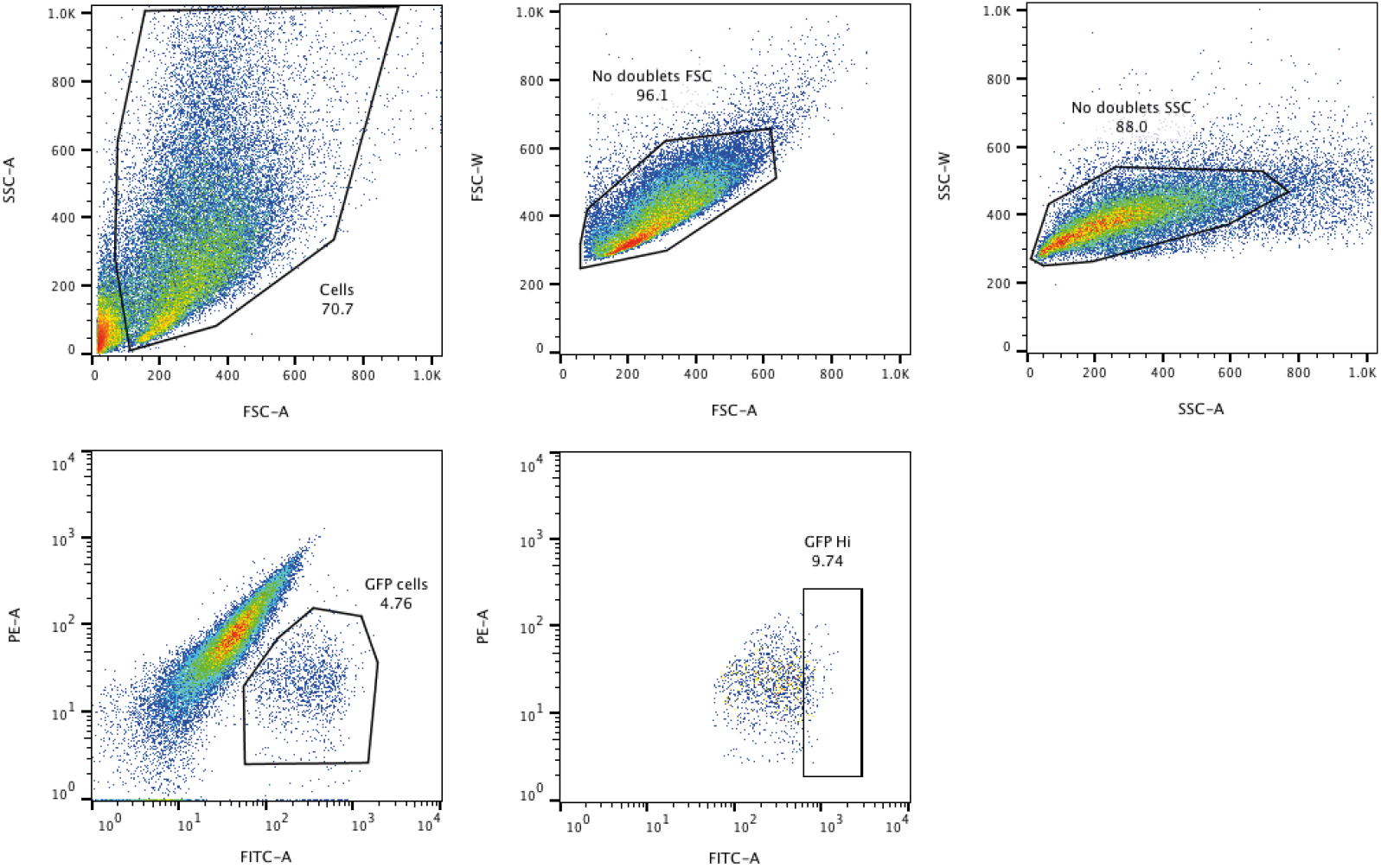
Isolation of satellite cells by FACS. Satellite cells were isolated by FACS by gating first on size and granulosity (‘Cells’ gate), excluding doublets (‘No doublets FSC’ then ‘No doublets SSC’ gates) and gating on the GFP+ population (‘GFP cells’ gate). Pax7-nGFP^Hi^ cells (top 10% highest nGFP-expressing cells, ‘GFP Hi’ gate) were sorted.

**Supplementary Table 1.**
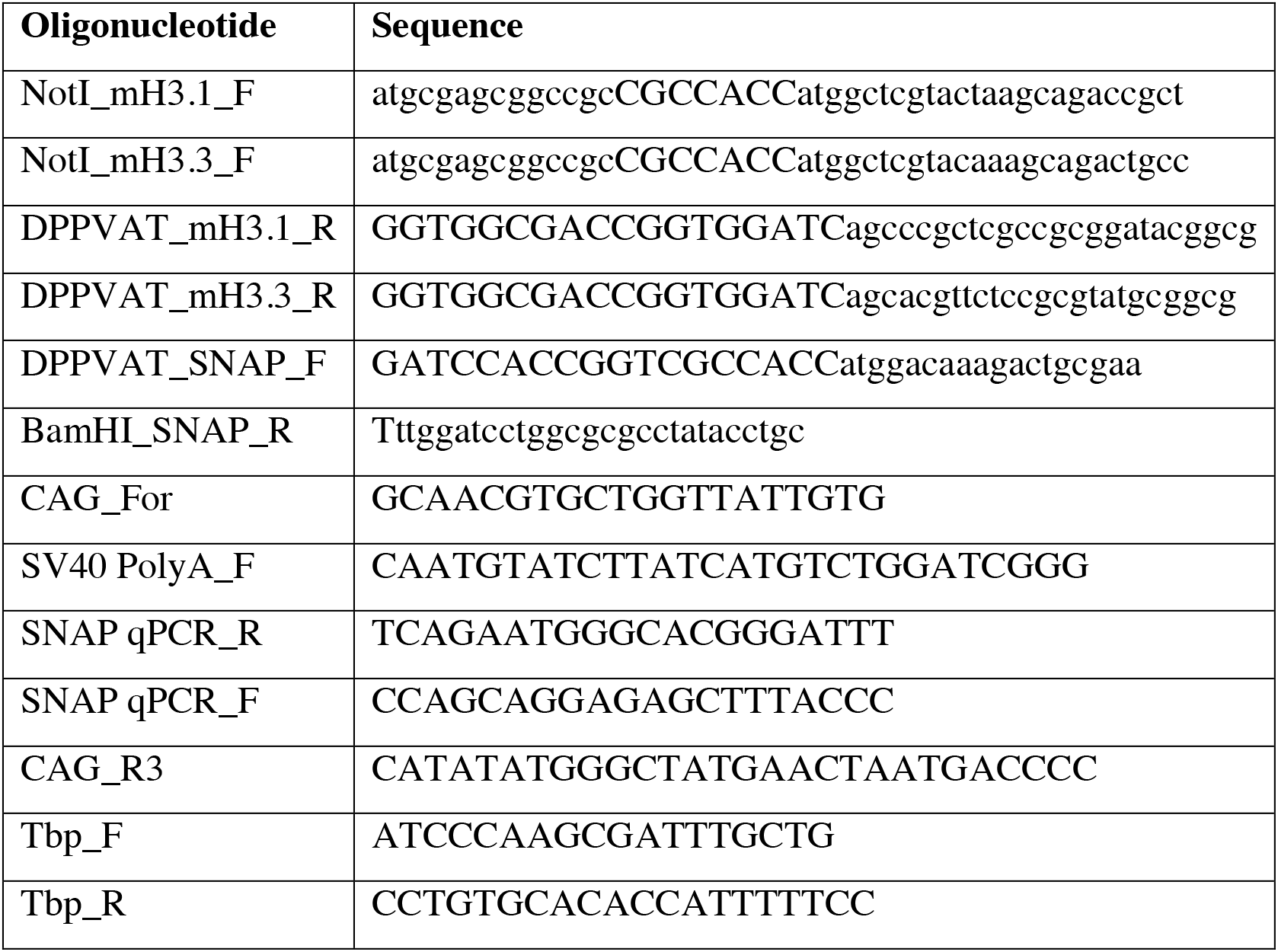
Oligonucleotides used in this study.

## Supplementary movies

**Supplementary Movie 1**.

Videomicroscopy of a Pax7-nGFP^Hi^ cell isolated from TA muscle at 5 dpi and plated on an asymmetric micropattern (not visible), corresponding to Fig. 1c left. The movie was started 3 h after plating and continued for 40 h; images were taken every 9 min (bright-field, 10× magnification).

**Supplementary Movie 2**.

Videomicroscopy of a Pax7-nGFP^Hi^ cell isolated from TA muscle at 5 dpi and plated on an asymmetric micropattern (not visible), corresponding to Fig. 1c middle. The movie was started 3 h after plating and continued for 40 h; images were taken every 9 min (bright-field, 10× magnification).

**Supplementary Movie 3**.

Videomicroscopy of a Pax7-nGFP^Hi^ cell isolated from TA muscle at 5 dpi and plated on an asymmetric micropattern (not visible), corresponding to Fig. 1c right. The movie was started 3 h after plating and continued for 40 h; images were taken every 9 min (bright-field, 10× magnification).

